# Run or glide: muscles are indifferent while the tendon takes the strain

**DOI:** 10.64898/2026.05.15.725315

**Authors:** Øyvind Gløersen, Anders Lundervold, Amelie Werkhausen

## Abstract

Conventional diagonal stride skiing traditionally includes a glide phase, characterised by a period of relatively passive gliding on one ski. While the glide phase may take advantage of low ski-snow friction, it does not exhibit the same whole-cycle mechanical energy fluctuations seen in running or walking on foot. A new sub-technique, known as running style, substantially reduces the glide phase and may alter the role of elastic tissues, making the movement pattern more similar to uphill running on foot in its temporal organisation.

We examined knee extensor and plantar flexor muscle–tendon behaviour in eight competitive skiers performing conventional diagonal and running techniques on a treadmill inclined at 10°. Using synchronised ultrasonography, 3D kinematics, ski forces and EMG, we quantified gastrocnemius medialis and vastus lateralis fascicle and muscle-tendon unit (MTU) dynamics in both the running (RUN) and conventional (CON) styles.

Shorter glide and total cycle durations during RUN shifted MTU peak length and velocity earlier during the kick phase. Fascicles in both muscles operated at similar velocities across techniques, showing MTU–fascicle decoupling. Vastus lateralis fascicles shortened at higher absolute peak velocities than gastrocnemius in both conditions, while normalised velocities were similar. RUN increased preactivation and advanced EMG timing, while integrated EMG during the kick was lower compared to CON.

These findings suggest that, despite large shifts in external mechanics between glide-based and more running-like skiing, elastic tissues may help stabilise fascicle behaviour and preserve a similar contractile strategy across muscles and techniques.

## Introduction

During walking and running on foot, elastic structures in the muscle-tendon complex allow efficient muscle work (1). For lower limb muscles, elastic strain may be reutilised so that muscles can reach high force potential by operating at favourable lengths and velocities, with surprisingly small differences between lower and upper leg muscles (2, 3) despite the differences in tendon properties (4). Muscle activity is greatest during the support phase, and running in particular is characterised by a clear preactivation in lower and upper limb muscles (5).

Studying how locomotion strategies are affected by manipulating the external conditions can be useful to understand the underlying mechanisms for efficient locomotion. For instance, conditions that require positive net work, such as running at an incline or accelerating, have greater muscle fascicle shortening than level or constant speed running (1, 6). Power-glide locomotion, in which active propulsion alternates with relatively passive glide periods, is well known in aerial and aquatic species and can also be enabled terrestrially through tools such as wheels, skis or skates (7). Locomotion over ice or snow is therefore another manipulation, where humans exploit low gliding friction to achieve efficient tool-assisted terrestrial movement (8, 9). This suggests that muscle–tendon coordination strategies established during upright bipedal locomotion may generalise to externally constrained, tool-assisted movement patterns.

Cross-country skiing, which is typically performed over undulating terrain, including both up- and downhills, includes the use of poles, making it a four-limbed locomotion mode. The conventional diagonal stride technique (CON) is a distinct movement strategy in classical cross-country skiing, typically used in uphill terrain. CON consists of a glide, kick (or push-off) and swing phase, or when performed on roller skis, a rolling phase, ski stop and swing phase (10, 11) (Figure 1A). The glide phase is non-propulsive, making CON a terrestrial power-glide gait rather than standard running. Although diagonal stride has been described as running-like, or as a form of grounded running, in some respects (12), the glide phase makes the mechanical energy fluctuations of CON distinct from running on foot (7). This is important because it indicates that CON cannot rely on the same whole-cycle mechanical energy exchange as terrestrial running, which we define here as non-assisted running on foot. Propulsion is mainly generated during the kick phase, which requires static friction between the ski and the surface (13). The static friction comes from grip wax under the mid section of the ski being pushed into contact with the snow. Hence, ensuring sufficient static friction requires an explosive movement at the kick phase onset. Since this explosive kick follows the non-propulsive glide phase, CON likely relies less on whole-cycle energy exchange and more on active propulsion than terrestrial running (7, 14).

**Fig. 1.**
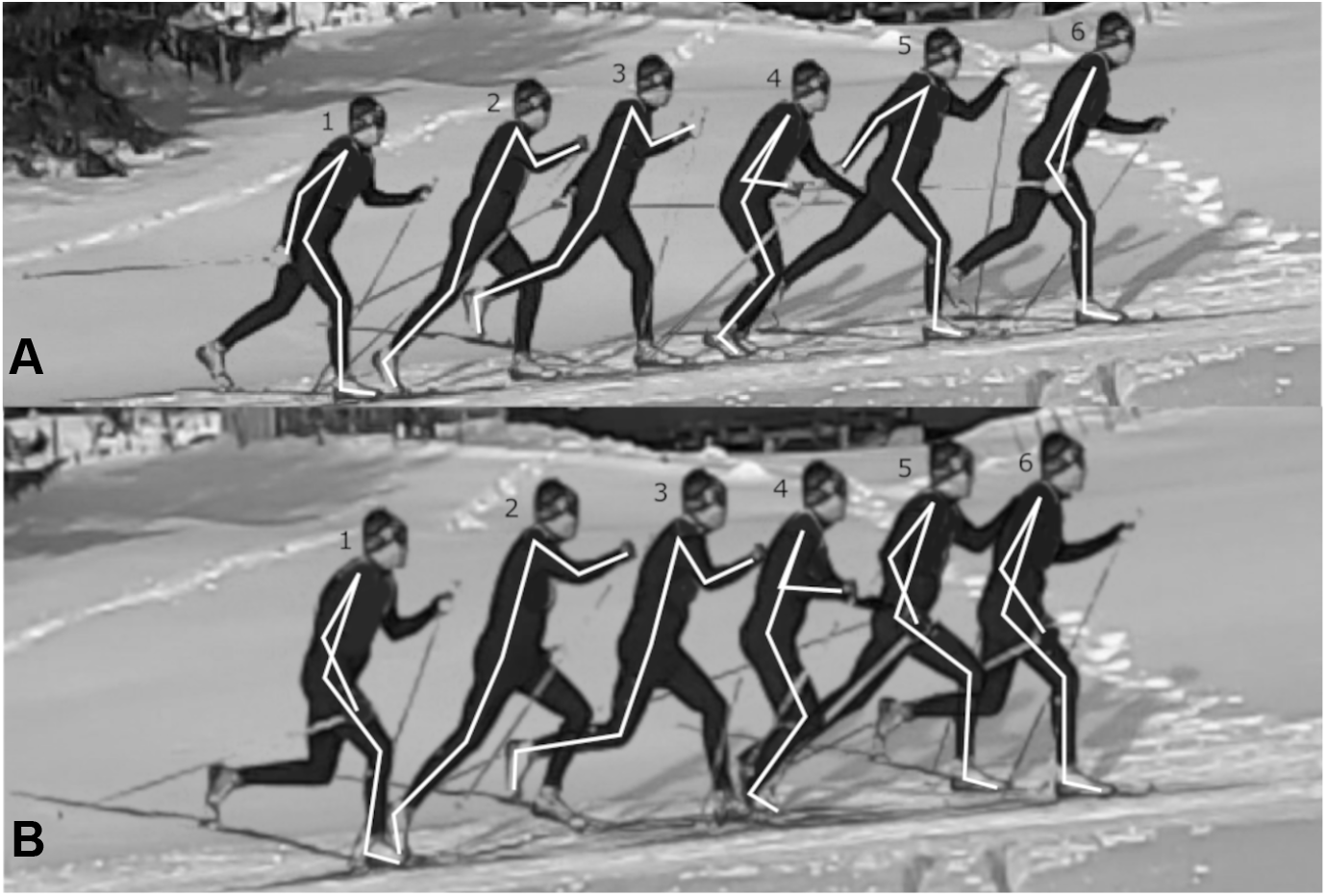
A full cycle of the two different styles of diagonal stride skiing. **A**: conventional style with a glide phase (CON). For the right leg, the poses show: start kick (1), start swing (2), mid-swing (3), start glide (4), mid-glide (5) and start kick (6). **B**: running style without a glide phase (RUN). For the right leg, the poses show: start kick (1), start swing (2), swing phase (3, 4 and 5) and start kick (6). One of the authors (ØG) is shown in this image, with consent for publication.

Komi and Norman were the first to describe a stretch-shortening cycle during diagonal stride skiing with a proximal-to-distal joint sequence and activation pattern (15). Recently, results from our group confirmed that plantar flexor fascicle kinematics were decoupled from limb kinematics during the kick phase, resulting in an MTU stretch-shortening pattern while fascicles shorten during the kick phase (14). This mechanism takes advantage of a catapult-like action where active muscle shortening enables elastic energy storage in the tendon that can be released at high rates (16, 17). Importantly, this catapult-like action is initiated by active muscle work, following a power amplification strategy, which differs from level and constant velocity running on foot, where the stretch-shortening cycle contributes more strongly to energy conservation (18).

Running style diagonal stride (RUN) is a distinct sub-technique in cross-country skiing that more closely resembles terrestrial running in its temporal organisation, albeit performed on skis. Unlike the conventional style (CON), RUN has an aerial phase and lacks, or has a very short, glide phase (Figure 1B). RUN also has a shorter cycle duration (approximately 28 %), earlier peak kick force, and greater knee joint excursion compared to CON (10).

Whereas previous work has distinguished glide-based and more running-like whole-body mechanics in skiing (7, 10), less is known about how these differences are expressed in muscle and tendon behaviour, especially across lower and upper leg muscles. In this study, we compare muscle and MTU behaviour in RUN and CON in both a plantar flexor (gastrocnemius medialis) and a knee extensor (vastus lateralis) muscle. This provides insight into whether running-like and power-glide strategies are reflected in force and energy transmission at two key joints (ankle and knee), while also allowing comparison of MTUs with different tendon properties (i.e., the highly compliant Achilles tendon and the stiffer patellar tendon).

We hypothesise that RUN, compared to CON, will show greater MTU length changes and velocities during the stretch-shortening cycle, whereas fascicles operate at similar velocities in both conditions owing to modulation by elastic tissue. If fascicle behaviour is preserved despite altered MTU mechanics, the elastic element (EE) should accommodate the difference, with greater EE length changes in the gastrocnemius medialis where the Achilles tendon is more compliant than the patellar tendon. Based on findings from terrestrial walking and running, we expect similar fascicle velocities in the vastus lateralis and the gastrocnemius in both RUN and CON. Given the 10° incline, we further expect greater fascicle shortening in both muscles than reported for level running, particularly for vastus lateralis, where fascicles are reported to operate near-isometrically during level running (3, 19). Furthermore, we hypothesise greater muscle preactivation at touchdown, as well as increased and earlier muscle activity during the kick in RUN compared to CON, in both muscles due to an earlier peak ski force during the kick.

## Methods

### Participants

Eight competitive cross-country skiers (4 female and 4 male; age: 27 ± 5 yr; height: 177 ± 9 cm; body mass: 72 ± 7 kg) were included in the study. Each participant had at least 5 yr of racing experience and prior experience roller skiing on a treadmill, and all provided written consent. The present analyses are based on the same data collection as Werkhausen et al. (14), but include the RUN condition and the vastus lateralis, which were not previously analysed. The study was approved by the institutional ethics board at the Norwegian School of Sport Sciences (approval 137-180620).

### Experimental protocol

Before testing, participants completed a 10 min low-intensity warm-up on a motorised treadmill (size 3.0 × 4.5 m, Rodby, Vänge, Sweden). To better reproduce on-snow conditions, the front and rear wheels of the roller skis (IDT Sport classic, Lena, Norway) were swapped so that the ratchet wheel was positioned at the front. Swix Triac 3.0 ski poles (BRAV Norge, Norway) fitted with treadmill ferrules were used throughout testing.

Testing was conducted on the same motorised treadmill, with a speed of 2.5 m · s^−1^ and an incline of 10°. Two conditions were tested: conventional-style diagonal stride (CON; similar to the sequence shown in Figure 1A) and running-style diagonal stride (RUN; similar to the sequence shown in Figure 1B). All participants were familiar with both styles and were instructed by the test leader which style to use.

Data were recorded for 10 s, with two trials collected for each condition. Participants rested for 2 min between trials.

### Joint kinematics and muscle–tendon unit length

Three-dimensional kinematics were collected at 150 Hz with a 12-camera motion capture system (Qualisys 400/700-series, Qualisys AB, Gothenburg, Sweden) using 26 reflective markers (12 mm). Twelve markers were placed on the pelvis and right leg at anatomical landmarks. Hip joint centres were defined from the anterior and posterior superior iliac spines (20). The knee and ankle joint centres were defined as the midpoints between the femoral epicondyles and the malleoli, respectively.

The foot segment was represented by boot-mounted markers at the calcaneus and the first, second, and fifth metatarsal heads. Three-marker clusters were used to represent the thigh and shank segments. Joint angles at the ankle, knee, and hip were computed using the approach described in (21). An additional eight markers were attached to the roller skis and poles. Reference frames were established from a static standing trial. Marker trajectories were smoothed with a zerophase second-order Butterworth low-pass filter (15 Hz).

Gastrocnemius medialis and vastus lateralis muscle-tendon unit (MTU) lengths were estimated from ankle and knee angles and scaled by shank and thigh length following (22). MTU velocity was obtained by differentiating the MTU length signal.

### Muscle fascicle length

Linear-array ultrasound was used to image gastrocnemius medialis and vastus lateralis fascicles. The probes (LV8-5N60-A2 ArtUs for gastrocnemius medialis and LogicScan 128 for vastus lateralis; Telemed, Vilnius, Lithuania) sampled at 117 Hz and 82 Hz, respectively, with a 50 mm field of view and 65 mm scan depth. Each probe was fixed to the skin with a custom holder and tape. We used a fully automated tracking algorithm (23) to estimate fascicle length and pennation angle. Pennation angle was defined as the angle between the fascicle and the deep aponeurosis. When a fascicle extended beyond the image boundaries, its path was linearly extrapolated. Both fascicle metrics were interpolated to 150 Hz and then low-pass filtered. Fascicle velocity was taken as the time derivative of fascicle length.

The length of the elastic element (EE), representing the tendinous and aponeurotic tissues in series with the fascicles, was estimated as

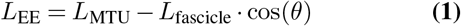

where *L*_MTU_ is the muscle-tendon unit length, *L*_fascicle_ is the fascicle length, and *θ* is the pennation angle. EE velocity was obtained by differentiating the EE length signal.

All tracking files were inspected independently by two of the authors, who then reached consensus on inclusion or exclusion from further analysis. Only participants with acceptable tracking in both muscles and both conditions were included, which led to the exclusion of five of the original 13 enrolled participants. In all five excluded participants, vastus lateralis image quality was too poor for acceptable fascicle tracking.

### Forces under the ski bindings

Vertical forces beneath the ski binding were estimated with two force-sensing resistors (Flexiforce A201, Tekscan, 0-445 N, 2100 Hz) mounted under the binding, as described in (14).

### Muscle activity

EMG signals were recorded at 2100 Hz with an Aktos Myon system. After skin preparation, bipolar surface electrodes were positioned over gastrocnemius lateralis and vastus lateralis in accordance with SENIAM (24). EMG was not recorded from medial gastrocnemius because the ultrasound probe occupied that site. The signals were band-pass filtered (10 Hz to 500 Hz) and converted to RMS using a moving 100 ms window. RMS amplitudes were normalised to the participant-specific maximum observed during CON.

### Cycle characteristics

A cycle spanned the interval between successive kick onsets. Kick onset was identified when ski velocity matched treadmill velocity within 0.5 km · h^−1^ (25). Ski position was obtained from the front wheel marker, and ski velocity was calculated as its time derivative. The coordinate system was rotated so that one axis aligned with the treadmill direction. This velocity signal was filtered with a bidirectional second-order Butterworth filter (20 Hz).

Each cycle was divided into kick, swing, and glide phases. Swing onset was defined as the point where ski velocity exceeded treadmill velocity by 0.5 km · h^−1^. Glide onset corresponded to the instant of maximal increase in force-sensing resistor (FSR) force (peak slope). In addition to the kick, swing and glide phases, we defined an *extended kick phase*, which also included a short period prior to the kick. Specifically, the original kick phase was elongated by 30 % into the preceding glide phase, corresponding to 70 ms to 80 ms. The extended kick phase was used for statistical analyses related to stretch shortening behaviour.

### Data Analysis

Five cycles per participant and condition were entered into the analysis. Cycles that deviated more than three scaled median absolute deviations from each participant’s median cycle were detected and removed using dynamic time warping (26). All data were time-normalised to 101 points per cycle. Scalar outcomes extracted for the kick phase included mean and peak fascicle and MTU length changes, mean and peak velocities, mean normalised EMG, and iEMG. Negative velocities denote shortening, whereas positive velocities denote lengthening.

### Statistics

#### Dependent variable definitions

To test the hypothesis that MTU length changes and velocities were higher in RUN than in CON during the kick phase, we calculated peak MTU stretch (difference from the local maximum during the kick to the local minimum before the kick) and peak MTU shortening (difference from the local maximum during the kick to the local minimum after the kick).

Fascicle contraction was defined as the difference between fascicle length at kick onset and the minimum fascicle length during the kick phase, reported in absolute units (cm) and normalised to fascicle length at kick onset (L_0_). To examine whether fascicle contraction velocities differed between conditions (RUN vs CON) and muscles (vastus lateralis and gastrocnemius medialis), peak fascicle contraction velocity was determined using a 100 ms moving average to reduce differentiation noise, and mean contraction velocity was calculated across the kick phase. Since fascicle length differed between muscles, we analysed both absolute velocities (cm · s^−1^) and velocities normalised to fascicle length at kick onset (L_0_ · s^−1^). The normalised velocities were used for inferences on muscle contractile conditions, since they were assumed to be approximately proportional to sarcomere contraction velocity, while absolute velocities were used for inferences on decoupling between muscle and MTU.

To assess differences in muscle activation, we computed EMG amplitude at kick phase onset (preactivation), time to peak EMG relative to kick phase onset, and integrated EMG (iEMG) over the kick phase. We also computed peak ski forces, time to peak ski force relative to kick phase onset, and the time integral of net ski force over the kick phase.

#### Statistical inference

Separate two-way repeated-measures ANOVAs were performed for each dependent variable described above, with condition (RUN and CON) and muscle (vastus lateralis or gastrocnemius medialis/lateralis, depending on the measurement modality) as factors. Simple effects with Tukey-Cramer multiple comparison corrections were calculated if the interaction between condition and muscle was significant. Both condition by muscle and muscle by condition interactions were calculated. Normality of the marginal differences was assessed using the Lilliefors test and graphical inspection of quantile-quantile plots.

For ski forces, we compared peak ski force, time to peak ski force, and the force-time integral between RUN and CON using paired *t*-tests. Paired *t*-tests were also used to compare cycle and phase durations (kick, swing and glide) between RUN and CON.

Lastly, we performed paired *t*-tests over the extended kick phase (see the Cycle characteristics subsection) for each dependent variable using statistical parametric mapping (SPM) (27).

All data and statistical analyses were conducted in Matlab R2024a. Statistical significance was set to *α* = 0.05.

## Results

### Temporal parameters

The duration of the whole skiing cycle and its parts, i.e., kick, swing and glide, are presented in Table 1. RUN had 0.35 s shorter cycles compared to CON, whereas both the glide and kick phases were statistically shorter in RUN compared to CON. Swing duration did not differ between conditions.

**Table 1.**
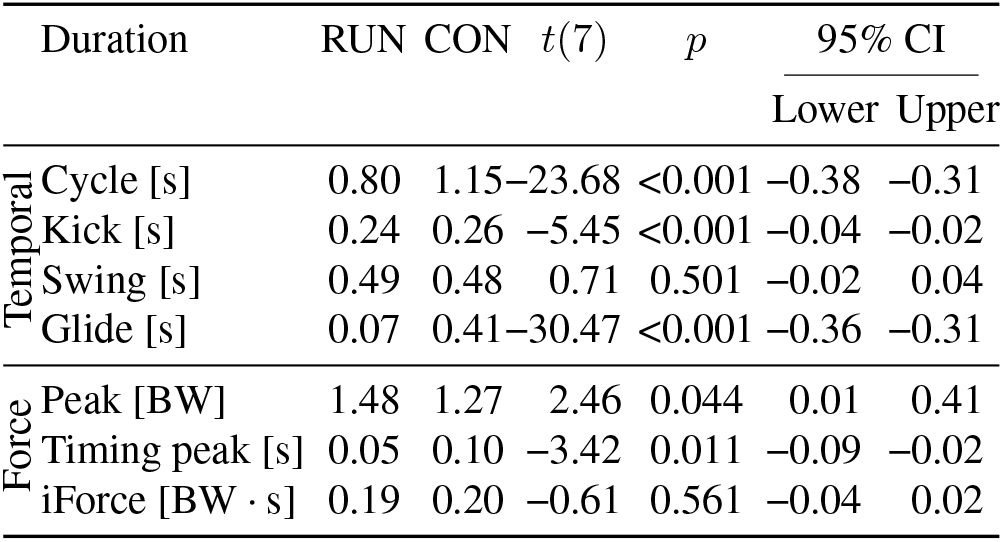
Temporal and binding normal force parameters during running on skis (RUN) and conventional diagonal stride (CON) (*n* = 8). Force variables are expressed relative to body weight (BW). Peak denotes peak normal force during the kick phase. Time to peak denotes the time from kick onset to peak force. iForce denotes the time integral of the normal ski force over the kick phase.

### Force

Peak ski force, time to peak force relative to kick onset, and force-time integral over the kick phase are presented in Table 1. Peak ski force was 16 % higher during RUN compared to CON, and occurred earlier in the kick phase (0.05 s vs 0.10 s after kick onset, respectively). There was no difference in the force-time integral over the kick phase; however, it should be noted that the kick onset definition used in this study does not include the initial ground contact period of RUN. The full force-time trace is presented in Figure 2B. The SPM analysis indicated significantly higher ski forces during RUN compared to CON from 6 % to 38 % of the extended kick cycle, corresponding to approximately 59 ms before kick onset to 47 ms after kick onset.

**Fig. 2.**
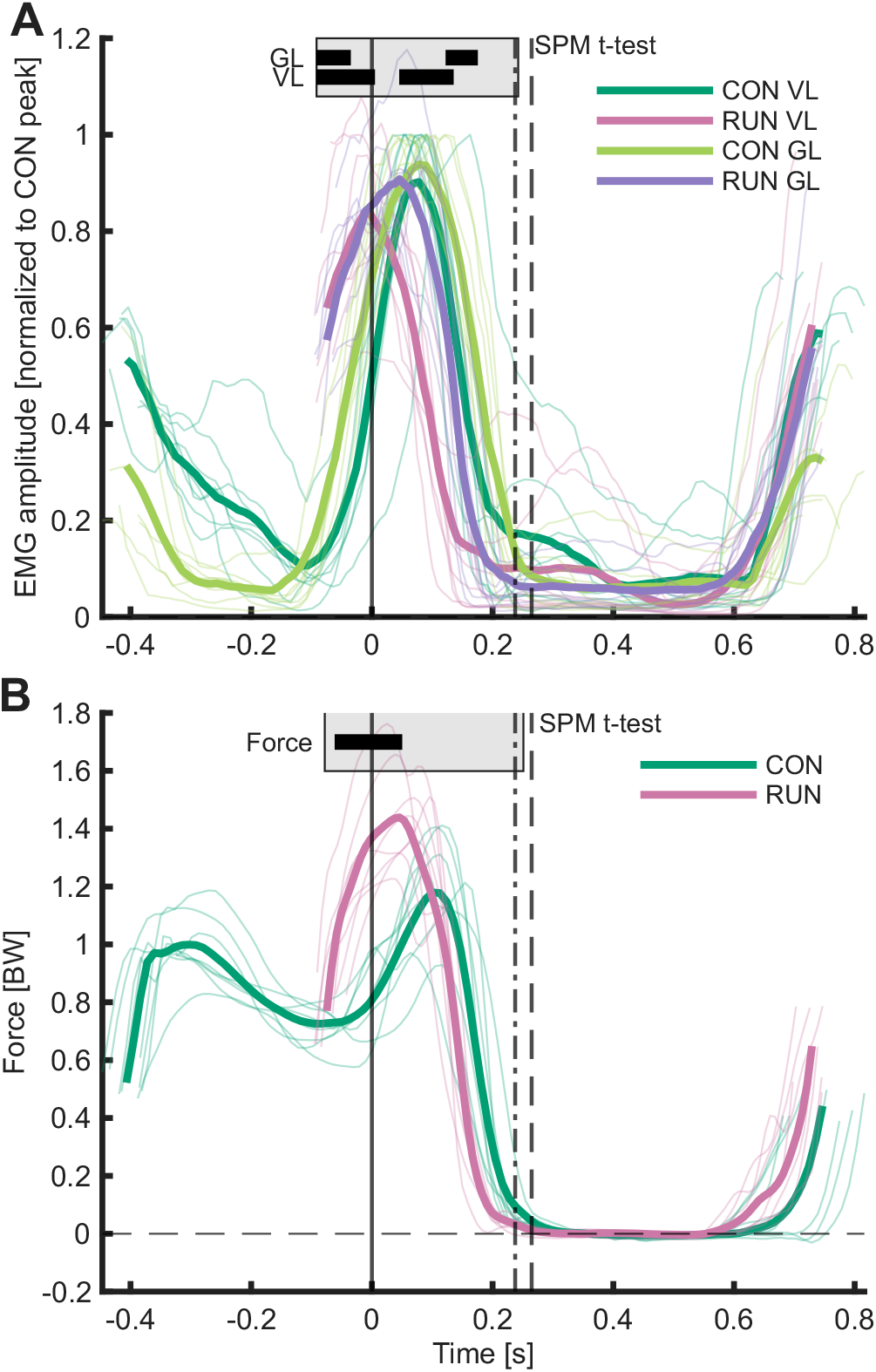
EMG activity in the vastus lateralis and gastrocnemius lateralis is shown in A, and ski force in B, during running (RUN) and conventional diagonal stride (CON). Thin lines indicate per-participant average cycles, and thick lines indicate the average across participants (*n* = 8). The solid vertical line at *t* = 0 marks the start of the kick phase, while the dotted and dashed lines mark the end of the kick phase for RUN and CON, respectively. The gray box, covering the “extended kick phase” (see Methods section for definition) and hence also covering a time period directly prior to the kick, shows the result of the *t*-test between RUN and CON using statistical parametric mapping. A thick solid black line indicates a significant difference between RUN and CON. Note that this graphical representation of the statistical tests is approximate, since the kick phase duration differed slightly between CON and RUN. See the manuscript body for exact percentages where the extended kick phase differed significantly between RUN and CON.

### MTU kinematics

Vastus lateralis and gastrocnemius medialis MTUs showed a stretch-shortening pattern during the kick phase of RUN and CON (Figure 3A,B). For gastrocnemius medialis, MTU shortening started earlier relative to kick onset in RUN compared to CON. This is evident in the SPM comparison, showing higher contraction velocities (Figure 3B) and shorter MTU lengths (Figure 3A) in the middle of the kick phase. MTU shortening also started earlier in RUN compared to CON for vastus lateralis, as indicated by the higher contraction velocities during the first half of the kick phase in the SPM analysis (Figure 3B). Furthermore, vastus lateralis MTU was longer in RUN compared to CON around the start of the kick phase (Figure 3A).

**Fig. 3.**
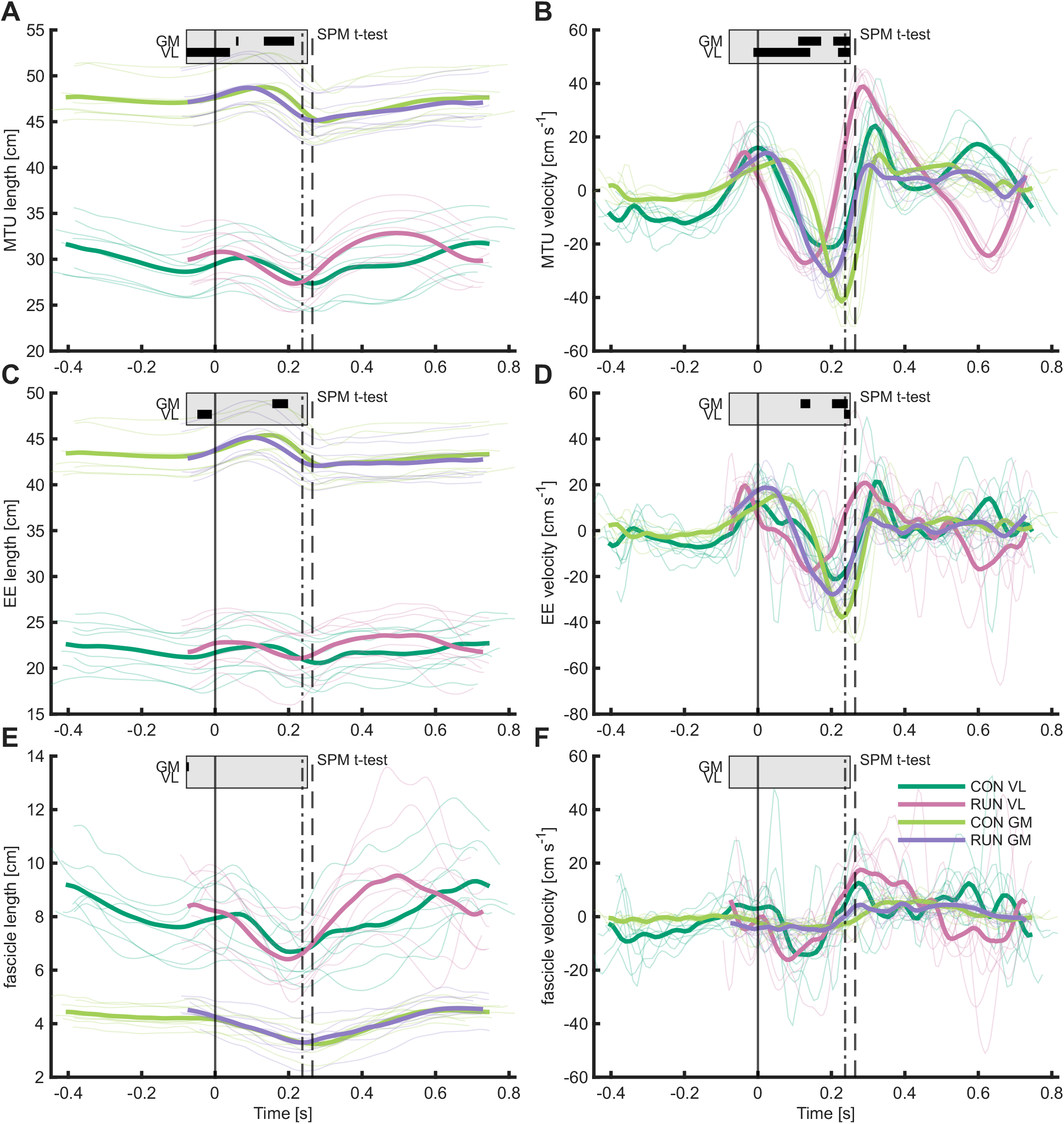
Mean MTU (muscle-tendon unit) length and velocity are shown in A and B, respectively, mean EE (elastic element) length and velocity in C and D, respectively, and mean fascicle length and velocity in E and F, respectively, for the vastus lateralis and gastrocnemius medialis during running (RUN) and conventional diagonal stride (CON). Thin lines indicate per-participant average cycles, and thick lines indicate the average across participants (*n* = 8). The solid vertical line at *t* = 0 marks the start of the kick phase, while the dotted and dashed lines mark the end of the kick phase for RUN and CON, respectively. The gray box, covering the “extended kick phase” (see Methods section for definition) and hence also covering a time period directly prior to the kick, shows the result of the *t*-test between RUN and CON using statistical parametric mapping. A thick solid black line indicates a significant difference in length or velocity between the RUN and CON conditions. Note that this graphical representation of the statistical tests is approximate, since the kick phase duration differed slightly between CON and RUN. See the manuscript body for exact percentages where the extended kick phase differed significantly between RUN and CON.

We found interaction effects in the ANOVA for MTU lengthening (*F* (1, 7) = 9.07, *p* = 0.020) and shortening (*F* (1, 7) = 20.15, *p* = 0.003). Simple effects for condition by muscle indicated no differences in lengthening between RUN and CON (*p* = 0.063 for vastus lateralis and *p* = 0.846 for gastrocnemius medialis). Comparing MTU by condition, gastrocnemius lengthening was greater than vastus lateralis lengthening during RUN (*p* = 0.015), while there was no difference during CON (*p* = 0.952). Gastrocnemius MTU shortened less in RUN compared to CON (*p* = 0.048), while vastus lateralis tended to shorten more (*p* = 0.050). Comparing MTU by condition, gastrocnemius peak shortening was greater than vastus lateralis shortening during CON (*p* = 0.011), while there was no difference during RUN (*p* = 0.480). Average values for both conditions and muscles are shown in Table 2.

**Table 2.** Muscle-tendon unit (MTU), elastic element (EE), fascicle and EMG parameters during running on skis (RUN) and diagonal stride (CON) in the vastus lateralis (VL) and gastrocnemius (GM). MTU and fascicle parameters for the gastrocnemius are measured from the medial head, whereas EMG activity is measured from the lateral head (*n* = 8).

| Variable | Parameter | RUN |  |  |  | CON |  |  |  | Simple effects |  |  |  |
| --- | --- | --- | --- | --- | --- | --- | --- | --- | --- | --- | --- | --- | --- |
|  |  | VL |  | GM |  | VL |  | GM |  | Condition by muscle |  | Muscle by condition |  |
| | | Mean | SD | Mean | SD | Mean | SD | Mean | SD | $p(C \vert VL)$ | $p(C \vert GM)$ | $p(M \vert RUN)$ | $p(M \vert CON)$ |
| MTU | Lengthening [cm]<sup>†</sup> | 1.18 | 0.33 | 1.83 | 0.37 | 1.87 | 0.78 | 1.85 | 0.46 | 0.063 | 0.846 | 0.015 | 0.952 |
|  | Shortening [cm]<sup>†</sup> | 3.59 | 0.39 | 3.82 | 0.66 | 3.09 | 0.64 | 4.14 | 0.46 | 0.050 | 0.048 | 0.480 | 0.011 |
|  | Peak velocity [cm · s<sup>-1</sup>]<sup>†</sup> | -25.73 | 2.48 | -29.47 | 4.15 | -21.21 | 5.11 | -36.05 | 3.04 | 0.009 | <0.001 | 0.121 | <0.001 |
| EE | Lengthening [cm]<sup>‡</sup> | 1.89 | 0.63 | 2.63 | 0.42 | 1.89 | 0.86 | 2.56 | 0.49 | – | – | – | – |
|  | Shortening [cm]<sup>†</sup> | 2.03 | 1.21 | 3.33 | 0.70 | 1.42 | 1.16 | 3.70 | 0.54 | 0.150 | 0.035 | 0.052 | 0.003 |
| Fascicle | Mean velocity [cm · s<sup>-1</sup>] | -6.15 | 5.68 | -4.04 | 1.42 | -3.72 | 2.85 | -3.39 | 1.53 | – | – | – | – |
|  | Peak velocity [cm · s<sup>-1</sup>]<sup>‡</sup> | -16.15 | 9.64 | -6.17 | 1.63 | -15.57 | 5.38 | -5.90 | 2.44 | – | – | – | – |
|  | Contraction [cm]<sup>‡</sup> | 1.95 | 1.35 | 0.98 | 0.29 | 1.79 | 0.53 | 0.87 | 0.31 | – | – | – | – |
|  | Mean velocity norm. [L<sub>0</sub> · s<sup>-1</sup>]<sup>‡</sup> | -0.82 | 0.82 | -0.99 | 0.35 | -0.46 | 0.32 | -0.83 | 0.38 | – | – | – | – |
|  | Peak velocity norm. [L<sub>0</sub> · s<sup>-1</sup>] | -2.09 | 1.45 | -1.51 | 0.40 | -1.95 | 0.72 | -1.43 | 0.55 | – | – | – | – |
|  | Contraction norm. [L<sub>0</sub>] | 0.25 | 0.19 | 0.24 | 0.07 | 0.22 | 0.05 | 0.21 | 0.08 | – | – | – | – |
| EMG | Start kick [a.u.]<sup>†</sup> | 0.82 | 0.13 | 0.84 | 0.10 | 0.55 | 0.18 | 0.77 | 0.10 | <0.001 | 0.174 | 0.677 | 0.006 |
|  | Timing peak [s]<sup>†</sup> | 0.00 | 0.02 | 0.05 | 0.03 | 0.07 | 0.03 | 0.08 | 0.03 | <0.001 | 0.001 | 0.002 | 0.681 |
|  | iEMG kick [a.u. · s]<sup>†</sup> | 0.08 | 0.02 | 0.12 | 0.02 | 0.14 | 0.01 | 0.17 | 0.02 | <0.001 | <0.001 | <0.001 | <0.001 |
|  | Cycle average [a.u.]<sup>†</sup> | 0.26 | 0.05 | 0.30 | 0.05 | 0.28 | 0.04 | 0.25 | 0.04 | 0.097 | 0.002 | 0.070 | 0.173 |

Significance symbols: <sup>†</sup> interaction (simple effects tested), \* condition, <sup>‡</sup> muscle ( $p < 0.05$ ). Simple-effect columns show adjusted post-hoc p-values; – = not tested.

There was also an interaction effect for peak shortening velocity of the MTU (*F* (1, 7) = 81.70, *p <* 0.001). Simple effects indicated that gastrocnemius shortened at a lower peak velocity during RUN compared to CON (*p <* 0.001), while vastus lateralis shortened at a higher peak velocity (*p* = 0.009). Comparing muscles within each condition, gastrocnemius shortened faster than vastus lateralis during CON (*p <* 0.001), with no difference during RUN (*p* = 0.121).

### Elastic element behaviour

The EE exhibited a stretch-shortening pattern during the kick phase in both muscles and conditions (Figure 3C,D). For gastrocnemius medialis, the SPM analysis showed that the EE reached peak stretch earlier in RUN compared to CON, resulting in shorter EE lengths during the last half of the kick phase (Figure 3C). Furthermore, peak EE shortening velocity was lower in RUN than CON (Figure 3D). For vastus lateralis, the EE also reached peak stretch earlier in RUN, with a significant SPM region around the start of the kick phase where the EE was longer in RUN compared to CON (Figure 3C). Unlike gastrocnemius medialis, there was no sustained difference in EE shortening velocity between conditions for vastus lateralis (Figure 3D). For EE lengthening, there was a main effect of muscle (*F* (1, 7) = 14.30, *p* = 0.007), with greater lengthening in gastrocnemius medialis than in vastus lateralis, but no interaction or condition effects. For EE shortening, there was an interaction effect (*F* (1, 7) = 5.73, *p* = 0.048). Simple effects for condition by muscle indicated that gastrocnemius medialis EE shortened less in RUN compared to CON (*p* = 0.035), with no difference for vastus lateralis (*p* = 0.150). Comparing muscles within each condition, gastrocnemius medialis EE shortening was greater than vastus lateralis during CON (*p* = 0.003), with a similar trend during RUN (*p* = 0.052).

### Muscle behaviour

Vastus lateralis and gastrocnemius medialis fascicles shortened during the kick phase of RUN and CON (Figure 3E,F). For gastrocnemius medialis, the SPM analysis showed a very short period with longer fascicles in RUN compared to CON prior to the kick phase (Figure 3E). Apart from that, the SPM analysis did not indicate any differences in either fascicle length or velocity in either muscle during the extended kick phase.

There were no significant interaction or main effects for absolute fascicle contraction velocities during the kick phase, but there was a main effect of muscle for normalised fascicle velocities (*F* (1, 7) = 6.11, *p* = 0.043), with gastrocnemius medialis contracting 0.27 L_0_ · s^−1^ faster than vastus lateralis. For peak fascicle velocity, there was a main effect of muscle for absolute velocity (*F* (1, 7) = 15.81, *p* = 0.005), with vastus lateralis contracting 9.8 cm · s^−1^ faster than gastrocnemius medialis, while there were no interaction or main effects for normalised fascicle velocities.

### Muscle activity

EMG activity was greatest during the kick phase in both conditions and muscles (Figure 2A). During CON, a smaller peak in activity was present at the beginning of the glide phase for both muscles. When comparing RUN to CON, peak muscle activity occurred earlier in the cycle for both muscles (Figure 2A). Means and standard deviations for EMG activity at the start of the kick, time to peak activity, and iEMG during the kick are shown in Table 2.

The ANOVA revealed an interaction effect of condition and muscle for EMG activity at the start of the kick phase (*F* (1, 7) = 13.36, *p* = 0.008). Simple effects revealed a higher EMG activity in vastus lateralis in RUN compared to CON (*p <* 0.001), but no difference in gastrocnemius lateralis (*p* = 0.174). For muscle by condition comparisons, simple effects revealed a higher EMG activity in the gastrocnemius than in vastus lateralis during CON (*p* = 0.006), but no difference during RUN (*p* = 0.677).

There was an interaction effect for timing of peak EMG activity (*F* (1, 7) = 18.60, *p* = 0.004). Simple effects compar-ing RUN versus CON in each muscle showed that peak EMG occurred 29 ms earlier in gastrocnemius lateralis (*p* = 0.001) and 77 ms earlier in vastus lateralis (*p <* 0.001). Comparing muscles within each condition revealed that peak activity occurred 51 ms earlier in vastus lateralis than in gastrocnemius lateralis during RUN (*p* = 0.002), with no difference between muscles in CON (*p* = 0.681).

The ANOVA also showed a significant interaction effect for iEMG over the kick phase (*F* (1, 7) = 6.26, *p* = 0.041). Simple effects for condition within each muscle and for muscle within each condition were all significant (all *p <* 0.001), with iEMG lower for vastus lateralis than for gastrocnemius lateralis, and lower for RUN than CON. There was also a significant interaction effect for EMG averaged over one cycle (*F* (1, 7) = 50.47, *p <* 0.001). Simple effects showed that in gastrocnemius lateralis, average activity was higher in RUN compared to CON (*p* = 0.002), while there were no significant differences in vastus lateralis or the muscle by condition analysis.

## Discussion

This study compared MTU kinematics, muscle behaviour and muscle activity of the vastus lateralis and the gastrocnemius between two mechanically distinct styles of diagonal stride skiing: CON, which has a distinct gliding phase and represents a power-glide strategy, and RUN, which has a very short glide phase and largely resembles running on skis. Our main finding was that fascicle behaviour was largely similar in RUN and CON, despite clear differences in whole-cycle organisation, cycle timing and MTU dynamics. This suggests that elastic tissues may decouple muscle fascicle behaviour from whole-limb mechanics across distinct locomotor strategies on skis. The uncoupling between fascicle and MTU kinematics was evident in both muscles, but was more pronounced in gastrocnemius medialis than in vastus lateralis. Specifically, the ratio of MTU peak velocity to fascicle peak velocity was about 5.5 in gastrocnemius medialis versus 1.5 in vastus lateralis. Lastly, we found an earlier peak muscle activity in RUN, whereas greater EMG activity at kick onset was only observed for the vastus lateralis.

### RUN shifts the timing of the kick sequence

The shorter cycle duration in RUN compared to CON is mainly explained by a significantly reduced glide time during RUN. These results align with Pellegrini et al. (10), although our participants had longer cycle durations in both conditions. A similar incline of 10° was used in both studies; however, we used a speed of 9 km · h^−1^ and included both male and female skiers, while Pellegrini et al. used a speed of 10 km · h^−1^ for male skiers only. The differences in cycle and phase duration are generally in line with other studies showing shorter cycle duration with increasing speeds (e.g. (28)). Similar to our study, Pellegrini et al. (10) found no difference in swing phase during RUN or CON. Pellegrini et al. also found no difference in kick phase duration between conditions, whilst we found that RUN had a slightly but significantly 20 ms shorter kick phase than CON.

The shorter glide phase in RUN was accompanied by an earlier peak ski force and earlier MTU length and velocity changes in both muscles. For gastrocnemius medialis and vastus lateralis, MTU shortening started earlier relative to kick onset in RUN compared to CON. These results suggest that RUN is not merely CON with a shorter glide, but a technique in which the whole propulsive sequence is shifted earlier. This is in line with the description of RUN as a more running-like variant of diagonal stride (10). Whereas conventional diagonal stride has been described as a type of grounded running with the flight phase replaced by a glide phase (7, 12), the reduced glide and earlier kick events in RUN appear to move the technique closer to terrestrial uphill running in temporal organisation, rather than implying level-running-style spring-mass mechanics.

### Fascicle behaviour is preserved despite altered MTU mechanics

The fundamental reduction in glide time in RUN may impose differences in MTU and muscle behaviour.For both MTUs, we found an earlier peak length during RUN combined with greater joint range of motion as previously reported (10). The condition effect at the MTU level was muscle-specific, with lower peak shortening velocity in gastrocnemius medialis during RUN compared to CON, while vastus lateralis shortened at a higher peak velocity in RUN. Despite these differences in MTU mechanics, fascicle behaviour was remarkably similar between conditions. During the propulsive kick phase, we found a decoupling of MTU and fascicle kinematics for both muscles and conditions. During the kick, MTU velocities exceed those of the fascicles, so that the muscle contractile pattern is energy efficient during the whole ground contact phase (29) using favourable velocities (18). Similar to a previous study, skiers in our study also had a proximal-to-distal joint sequence (15), and our comparison of the technique conditions within each muscle suggests similar strategies for the ankle and knee extension.

Here we refer to peak fascicle velocity rather than average velocity over the kick phase, since average fascicle velocity was affected by fascicle elongation starting before the kick phase ended (Figure 3F). We found no statistical differences in peak normalised fascicle velocity between conditions or muscles, which supports our hypothesis of no differences in contractile conditions, even though the power-glide and more running-like strategies have clear differences in segment kinematics (10) and mechanical energy fluctuations (7). Thus, our findings are broadly consistent with terrestrial walking and running literature showing that elastic tissues can decouple fascicle behaviour from whole-limb mechanics (2, 3).

The degree of uncoupling was, however, not identical between muscles. It appeared more pronounced in gastrocnemius medialis than in vastus lateralis, which may be consistent with the higher compliance of the Achilles tendon compared with the patellar tendon (4). This is supported by the calculation of EE: gastrocnemius medialis EE lengthening and shortening were both greater than in vastus lateralis, and the condition-dependent reduction in gastrocnemius EE shortening during RUN mirrored the lower MTU peak shortening velocity in that muscle. Together, these results indicate greater elastic decoupling at the distal MTU, whereas fascicle and MTU behaviour appear more closely linked at the knee. Even so, both muscles converged on similar peak normalised fascicle velocities across conditions, suggesting that similar contractile conditions may be achieved through different mechanical solutions.

Although fascicle contraction velocity was moderate to low (1.4 L_0_ · s^−1^ to 2.1 L_0_ · s^−1^), our findings show a distinct fascicle contraction of about 0.20 L_0_ to 0.25 L_0_ in both muscles and both conditions. However, vastus lateralis showed notably high inter-subject variability in RUN (SD = 0.19 L_0_), suggesting considerable individual differences in VL fascicle behaviour during the running-style technique. While the mean fascicle contraction is in agreement with previous studies on level running for the gastrocnemius medialis (19), it is not for the vastus lateralis, where previous work shows almost isometric muscle behaviour during the stance phase (3, 19). The greater tendon decoupling in gastrocnemius medialis may be one reason why its behaviour resembles level running more closely than that of vastus lateralis. We speculate that the discrepancy from level running is mainly due to the 10° incline used in this study. Running up an incline requires net positive work, and uphill running progressively deviates from a pure spring-mass rebound as slope increases (30). Even on more moderate uphill slopes, the maximum possible whole-body elastic energy exchange is reduced relative to level running, although not abolished (31). Under such conditions, proximal joints are likely required to contribute more positive work, which could increase vastus lateralis fascicle shortening. An additional possibility is that the work distribution between joints in the lower extremities shifts from distal (primarily ankle) during level running, to proximal (primarily hip) during inclined running (32). Although the hip joint is the main power producer during inclined running, bi-articular hip extensor muscles might transfer power produced by knee extensors (33, 34). Together, these factors might explain why we observe similar fascicle shortening as in level running in the gastrocnemius medialis, but substantially greater contractions for the vastus lateralis.

### Neuromuscular control is retimed in RUN

The earlier MTU peak length observed during RUN may help explain the earlier muscle activity observed in RUN while maintaining a similar contractile pattern across conditions. This is consistent with our findings of earlier peak muscle activity in both muscles in RUN compared to CON. Greater EMG activity at the start of the kick was only observed for vastus lateralis, whereas gastrocnemius lateralis showed no difference between conditions at kick onset. Thus, the neuromuscular changes in RUN appear to reflect a retiming of activation rather than a uniform increase in muscle excitation, which are in line the MTU changes. Despite the shift in activation timing, overall activation patterns during the different skiing techniques were less affected than what has been reported when comparing terrestrial walking and running, particularly in the vastus lateralis (3), but also for the gastrocnemius medialis (35).

Although we predicted greater activation during the kick phase in RUN, iEMG was lower compared to CON, which may partly be explained by the earlier peak activation in RUN. Since the vastus lateralis activation peak in RUN occurs at kick onset, and the gastrocnemius peak shortly after (Table 2), the high activation levels that build up prior to and around kick onset are not fully captured by the kick-phase iEMG window. When comparing average EMG activation over the full cycle, we found only a modest increase for gastrocnemius lateralis, and no difference for vastus lateralis. Together with the preserved fascicle velocities, this suggests that earlier activation in RUN may contribute to earlier loading of elastic tissues and to the maintenance of similar contractile conditions despite the altered external timing. This is broadly consistent with evidence from *in situ* experiments in turkeys showing that activation timing modulates the relative contribution of elastic elements versus active fascicle mechanisms in the gastrocnemius, though that work exam-ined energy absorption rather than propulsion (36).

### Different elastic roles in power-glide and running-like skiing

Kehler et al. reported that diagonal stride skiing, similar to CON in the current study, was characterised by inphase fluctuations of the centre of mass, resembling running on foot more than walking (7). However, a key distinction was that during diagonal stride skiing, the kinetic energy generated during the kick phase was dissipated to rolling resistance during the glide phase. This implies that CON, unlike terrestrial running, cannot rely on the same whole-cycle mechanical energy exchange to the same extent. Rather, much of the energy required for the next kick must be generated actively.

Our results are consistent with elastic tissue loading and recoil in both CON, as discussed in our previous work (14), and RUN. The important distinction may therefore not be whether elastic tissues are used, but how they are used. In CON, slow fascicle shortening can tighten the tendon while the MTU shortens much faster, which is consistent with a catapult-like mechanism and power amplification. The higher gastrocnemius EE shortening velocity observed during CON compared to RUN supports this interpretation, suggesting a more pronounced elastic recoil and thus stronger power amplification during the CON kick phase. This condition difference was not observed for vastus lateralis, which is consistent with the lower compliance of the patellar tendon limiting the scope for catapult-like behaviour at the knee. In RUN, the shorter glide and earlier force and MTU events suggest a more continuous propulsive sequence, but our 10° uphill condition does not imply a level-running-like spring-mass strategy. Studies of terrestrial uphill running show that whole-body mechanics become more asymmetric and net-positive-work dominated as slope increases, even if some elastic exchange remains (30, 31). RUN may therefore be better viewed as an uphill-running-like strategy in which earlier loading and local tendon decoupling help preserve fascicle behaviour, rather than as a pure spring-mass bounce. Hence, CON may be viewed as a more muscle-generated bounce with glide-separated propulsion, whereas RUN appears to organise propulsion earlier and more continuously within the cycle.

### Perspective

An important implication is that in the field the two techniques may be advantageous under different external conditions. When ski-snow friction is low enough for the glide phase to preserve forward motion effectively, and when the required external work rate is moderate, CON may be more economical because a passive glide period can reduce the fraction of the cycle spent actively generating propulsion. In contrast, when ski-snow friction becomes too high, the incline too steep, or power output too high, RUN may be advantageous because it minimises the passive interval and shifts force generation earlier in the cycle. This interpretation remains speculative, as the present study did not measure metabolic cost, tangential ski forces, or manipulate friction and incline systematically.

The earlier and higher peak ski force in RUN may also have practical implications for on-snow skiing, where grip depends on sufficient normal force to press the grip wax into the snow. However, normal force alone does not determine grip. In CON, peak tangential force occurs after peak normal force (37, 38), and whether the same temporal relationship holds in RUN remains unknown. Measurements of both normal and tangential ski forces during RUN are therefore needed before concluding that the higher normal force improves grip.

### Limitations

Several limitations should be considered when interpreting these results. First, five of the original 13 participants were excluded due to poor vastus lateralis ultrasound image quality, resulting in a small sample (*n* = 8) that limits statistical power and generalisability. Second, EE length was estimated indirectly as the difference between modelled MTU length and the fascicle projection onto the muscle line of action. This estimate therefore includes both tendon and aponeurosis and is sensitive to errors in both the MTU model and the ultrasound-based fascicle measurements. Third, testing was performed on roller skis on a treadmill at a single speed and incline. The extent to which these findings transfer to on-snow skiing at varying speeds, inclines and snow conditions remains to be established. Finally, we did not measure metabolic cost or tangential ski forces, which limits our ability to draw conclusions about the economy of each technique or the functional role of the higher normal forces observed during RUN.

## Conclusion

At the same treadmill speed, RUN and CON differed in the timing and MTU mechanics of the kick phase, with RUN showing earlier force and MTU events. Despite these differences, fascicle behaviour was largely preserved across techniques, suggesting that elastic tissues may decouple contractile conditions from whole-limb mechanics in both gastrocnemius and vastus lateralis. The uncoupling between MTU and fascicle behaviour was more pronounced in gastrocnemius than in vastus lateralis, consistent with greater elastic decoupling at the ankle than at the knee. Together, these findings suggest that elastic tissues likely contribute in both techniques, but may serve different roles in a power-glide strategy (CON) and a more running-like strategy on skis (RUN). CON appears more consistent with power amplification during the kick after a glide phase, whereas RUN appears to organise propulsion in a more continuous, uphill-running-like manner.

